# Cloning, expression and characterisation of antigen-specific recombinant bat immunoglobulin from the black flying fox (*Pteropus alecto*)

**DOI:** 10.64898/2025.12.06.692532

**Authors:** Jun Jet Hen, Ariel Isaacs, Benjamin Liang, Tony Schountz, Keith Chappell, Paul R. Young, Naphak Modhiran, Daniel Watterson

**Affiliations:** School of Chemistry and Molecular Biosciences, The University of Queensland, Brisbane, Australia; Australian Infectious Disease Research Centre, The University of Queensland, Brisbane, Australia; Department of Microbiology, Immunology and Pathology, Colorado State University, Colorado, United States of America; Australian Institute for Bioengineering and Nanotechnology, The University of Queensland, Brisbane, Australia

## Abstract

Bats are natural reservoirs of viruses that cause severe disease in livestock and humans. Recent high-profile spillover events have directed significant attention towards the interplay between zoonotic viruses and antiviral immunity inherent to bats. Studies have highlighted that bats could harbour some deadly viruses without exhibiting outward symptoms. Various hypotheses have been proposed on how bats coexist with viruses; this includes dampened inflammation and altered innate immunity. However, there is limited literature on the humoral immune response in bats due to the scarcity of bat-specific reagents. To address this knowledge gap, we employed recombinant antibody design techniques to generate antigen-specific recombinant bat antibodies. This strategy involves combining the paratope of well-characterised antiviral antibodies with the IgG1 constant region of the black flying fox (*Pteropus alecto*). Characterisation of recombinant bat antibodies have revealed that they display canonical features of mammalian IgG. Additionally, recombinant bat antibodies display a binding and neutralising profile akin to human antibody counterparts. This approach provides much needed diagnostic tools and novel reagents to accelerate research into bat antibody immunity.

## Introduction

Bats are mammals of the order Chiroptera and are broadly divided into suborders Yinpterochiroptera and Yangochiroptera^1^. The propensity of bats to host zoonotic viruses has been at the centre of much attention^2–4^. Henipaviruses, lyssaviruses, filoviruses and betacoronaviruses are examples of highly pathogenic viruses hosted by bats that cause severe disease in humans and livestock^5–9^. Numerous endogenous viral elements (EVE) have recently been identified in bat genomes, suggesting that bats have long been associated with viral infections^10,11^. Indeed, the most distinctive features of bats are their ability to host diverse pathogens with no observable signs of disease^12–15^.

Several theories as to how bats coexist with viruses have been proposed. Genomic and transcriptomic studies on bats have revealed that their immune systems are broadly similar to other mammals with highly expanded immune-related genes^10,16–18^. Virus tolerance via altered pathogen sensing, enhanced innate immunity, and reduced inflammation are some of the examples of unusual immune adaptations that have been reported in bats^19–23^. However, little is known about the adaptive response in bats in response to viral infection. Evidence of lymphoid organs, B and T lymphocytes, augmented major histocompatibility complex (MHC) and the presence of Fc receptors (FcR) have been previously described^18,24–28^

In bats, all canonical mammalian immunoglobulin isotypes (IgM, IgG, IgA, IgE, IgD) have been described via genomics and proteonomics but distribution varies with species^18,29–32^. Interestingly, some evidence suggests that the kappa light chain is lost in some species of bats, and bats may preferentially use lambda light chain^33,34^. IgG subclass in bats varies by species and ranges from one (*A. jamaicensis, C. perspicillata*) to five (*M. Lucifugus)*^29,31,32^. Interestingly, some studies have noted inconsistent seroconversion and diminished antibody response in bats^35–42^. Others have found bats are capable of mounting a successful humoral response with high levels of seroconversion in experimental infections^43–49^.

Expanded germline repertoire with atypical amino acid and glycan composition have been reported in bats^29,32,50^. These observations led researchers to propose that combinatorial diversity of naïve B-cells may be more important for bats in controlling viral infections as opposed to somatic hypermutation and affinity maturation^31,32,50–52^. However, this concept around bat immunity is understudied due to the lack of bat-specific reagents and tools^53^. To better understand bat immunoglobulins and address the shortage of bat-specific tools, we generated recombinant bat antibodies (ReBAs) with paratope from the well-characterised anti-Nipah virus (NiV) fusion (F) glycoprotein antibody, 5B3. Here, we characterised ReBAs with a series of immunoassays to confirm their biochemical properties as well as their capacity to bind the target antigen and neutralise virus. Altogether, the complementary tools developed in this study will help expand our understanding of how bat antibodies recognise viral antigens and enable research into bat antiviral immunity.

## Results

### Recombinant bat antibodies construct designs

To generate ReBAs, we *de novo* synthesised and constructed mammalian expression plasmids containing constant heavy chain (Hc) or kappa (κ) light chain (Lc) derived from IgG1 of black flying foxes (Figure 1A, B). A *Kpn*I restriction site was introduced at the N-terminal of Hc or Lc plasmids for subsequent downstream cloning applications. To validate the ReBAs system, we selected two well-characterised antibodies as prototypes. These include anti-henipavirus F protein antibody, 5B3^54^, and the anti-influenza A hemagglutinin (HA) antibody, C05, that targets H1, H2 and H3 subtypes^55^. Both Hc and Lc variable domains from these antibodies were cloned in-frame with corresponding constant domains. Sequence alignment of the bat and human Hc constant domain shows that the bat constant region contains five (5) disulphide bonds whereas the human constant region contains six (6) disulphide bonds (Figure S1A). Specifically, the missing disulphide bond in the bat Hc region could be attributed to the deletion of the first cysteine at the hinge region (Figure 1C and S1). Bat Lc is similar to the human IgG1 Lcκ constant region; both have two interchain disulphide bonds and one C-terminal cysteine that enables intra-chain disulphide formation with the HC (Figure S1B). The glycan site position in the bat Hc region is similar to human IgG1 (Fig1D and S1). In addition, to assess the impact of the signal peptide on the recombinant bat antibodies expression, two N-terminal signal peptides, human signal peptide (Hsp) and bat signal peptide (Bsp) derived from black flying fox were incorporated into the antibody Hc. Sequence alignment shows 8 differences between Hsp and Bsp (Figure 1B). These features are summarised in Figures 1A and 1E, illustrating the envisaged configuration of black flying fox IgG1.

### Characterisation of recombinant bat antibodies

To characterise ReBAs or human IgG, we purified the cell culture supernatant using protein A (pA) or protein G (pG) resin columns. Figures 2A and 2B depict the SDS-PAGE analysis of the purified products under reducing conditions. Human IgG and recombinant bat antibodies exhibited similar band patterns, characterised by two prominent bands observed at approximately 50 and 25 kDa (Figures 2A and 2B). These bands correspond to typical mammalian antibody Hc (50kDa) and Lc (25kDa). Nevertheless, we note a small but discernable distinction between the molecular weights of the bat and human antibodies compared to human 5B3 (h5B3) and human C05 (hC05) with ReBAs (Figures 2A and 2B). The bat Lc consistently showed a smaller size than h5B3 and hC05 Lc (Figure 2A, lanes 2, 3, 5 and 6), whereas the bat Hc (Hsp and Bsp) of ReBAs appears marginally larger than that of h5B3 and hC05 (Figure 2A, lanes 2, 3, 5 and 6). This apparent shift in molecular weight may be attributed to differences in the isoelectric point (pI) between the proteins. Use of either human or bat signal peptides demonstrated no discernible differences, with bat 5B3Hsp (b5B3Hsp) and b5B3Bsp (Figure 2A, lanes 2 and 3) displaying similar band profiles and molecular weights. This result was consistent and also observed for bat C05Hsp (bC05Hsp) and bC05Bsp (Figure 2A, lanes 5 & 6). Similar profiles were observed with pG purified ReBAs (Figure 2B, lane 2, 3, 5 & 6). Still, some samples displayed faint bands outside the expected sizes of 50 and 25kDa (Figures 2A and B), suggesting the presence of impurities. Overall, the results obtained from SDS-PAGE analysis indicate that the recombinant bat antibodies – ReBAs IgG1, share a molecular weight profile similar to human IgG1 antibodies. This finding also shows that both pG and pA purification methods can effectively isolate ReBAs.

### Glycopeptidase F sensitivity analysis

To investigate the putative glycan sites within bat and human Hc (Figure 1D), we treated ReBAs with PNGase F, an enzyme that selectively removes N-linked oligosaccharides from proteins. For comparative analysis, h5B3 and bat b5B3 antibodies were examined under reducing conditions, with an untreated sample as the control. Upon treatment with PNGase F, the bands corresponding to Hc positioned at approximately 50kDa exhibited a downward shift compared to those of untreated samples (Figure 2E). This shift demonstrates that glycans were removed from the Hc, indicating that ReBAs are modified by complex N-linked glycosylation similar to human antibodies.

### Size-exclusion chromatography (SEC) and negative stain transmission electron microscopy (negTEM) of recombinant bat antibodies

To further evaluate the oligomeric state of bat antibodies, b5B3Hsp and b5B3Bsp antibodies were analysed by SEC on Superose 6 Increase 10/300 GL gel filtration column at physiological pH7.4. For both pA and pG purified antibodies, b5B3Hsp and b5B3Bsp yielded a peak at a retention volume of approximately 17.8mL (Figure 2C, D). Similarly, h5B3 eluted at a retention volume of about 17.8mL (Figure 2C, D). Substitution into the regression equation established using protein standards yielded a predicted molecular weight of approximately ∼145kDa (Figure S2), which closely resembles the molecular weight of human IgG1^56^. For pG purified antibodies, both b5B3Hsp and b5B3Bsp also eluted as a single peak at a retention volume of approximately 17.8mL (Figure 2D). These results show that the oligomeric state of ReBAs are similar to human IgG1. In addition, we also visualised the elution peak of SEC purified samples of b5B3Bsp with negative stain transmission electron microscopy (negTEM). The collected micrograph depicts particles of ∼10nm in size with distinct Y-shaped morphology (Figure 2F). These features are characteristic of a typical mammalian IgG and demonstrate that our construct yielded recombinant bat antibodies that resemble human IgG1.

### Recombinant bat 5B3 retains its specificity to NiV F protein

To determine if the antigen-binding functions of the ReBAs were preserved, we performed an indirect enzyme-linked immunosorbent assay (ELISA) against immobilised bat-borne NiV F protein and influenza virus HA protein with our panel of ReBAs. These antigens are stabilised by the Molecular Clamp (first-generation) and are described extensively elsewhere^57,58^. Due to the limited data available for anti-bat IgG secondary antibodies, we also tested a small panel of commercially available HRP conjugated anti-bat IgG secondary antibodies from Alpha Diagnostic, Bethyl and Novus Biologicals. After initial testing, we selected goat anti-bat antibodies from Novus Biologics (Cat: NB7238), that was raised against bat IgG Hc and Lc, for its favourable binding (lowest K_D_) to ReBAs (Figure S3). To further scrutinise this reagent, we supplemented our findings on ReBAs with an investigation into the reactivity of immunised bat sera to NiV F Clamp. Figure S4 shows that 2 out of 3 bats seroconverted 14 days after booster immmunisation with NiV F appended with foldon trimerisation domain, and NiV F specific antibodies were detected via ELISA with absorbance readings ranging from 1.2-1.3 absorbance units (AU). Conversely, those from control groups (N=3) have a lower absorbance (<0.5AU at the highest concentration) (Figure S4). Importantly, this indicates that the selected goat anti-bat secondary antibody could detect IgG from captive Jamaican fruit bats (*Artibeus jamaicensis*) and also ReBAs, which incorporates IgG constant regions of the black flying fox.

Next, we examined the binding kinetics of the henipavirus F-specific antibody, h5B3 and its bat derivatives that were purified with pA column. The pA purified b5B3Hsp and b5B3Bsp had apparent affinities of 0.89 and 0.72nM, respectively, which is within the nanomolar range of h5B3 with an apparent affinity (K_D_) of 0.51nM (Figure 3A, B). This analysis demonstrated that the apparents affinities of h5B3, b5B3Hsp and b5B3Bsp were comparable (Figure 3B). Interestingly, maximum specific binding (B_max_) values of h5B3 was higher at 2.79 AU, while the absorbance units of b5B3 were similar across Hsp and Bsp (1.52 - 1.83AU) (Figure 3B). pG purified b5B3 also exhibits a similar magnitude of binding (B_max_) but binding affinities were ∼two to five-fold lower compared to its pA purified human counterpart (Figure S5). As negative control, 5B3 ReBAs were used against HA antigens, and C05 ReBAs were tested on NiV F antigens to determine assay background levels. A trimerisation domain specific antibody, anti-Clamp1 (HIV1281), was used throughout as a positive control^59^. Overall, this shows that h5B3 and its bat derivative generally shares the same binding profile but subtle differences were observed in magnitude of binding (B_max_).

As a comparison, we also examined influenza HA specific hC05 and its bat derivatives against influenza (A/Brisbane/59/2007) HA proteins (Figures S6A and B). The apparent K_D_ values for pA purified hC05 was 0.24nM, while K_D_ values for bC05Hsp and bC05Bsp were 0.25 and 0.45nM respectively (Figure S6B). Interestingly, the binding affinity of hC05 to HA antigen was similar to bC05Hsp but is approximately two-fold higher than bC05Bsp. B_max_ values of hC05 pA was highest at 4.03AU, while readings for bC05Bsp and bC05Hsp were analogous at 3.38AU and 3.36AU (Figure S6B). Intriguingly, these results suggest that h5B3 amd hC05 may work marginally better than their bat derivatives. However, the comparison between human IgG and ReBAs should be interpreted cautiously due to the different secondary antibodies. Overall, this result confirms that chimeric ReBAs containing paratope retained their antigen-specific binding capacity.

### Recombinant bat antibody 5B3 neutralises NiV pseudovirus particles

To further investigate the neutralising capabilities of ReBAs, we used a NiV pseudovirus (NiV-pps) assay^58,60^. As illustrated in Figures 4A and 4B, b5B3Hsp and b5B3Bsp could neutralise NiV-pps well at a half-maximal inhibitory concentration (IC_50_) of 0.058nM and 0.084nM, while h5B3 has an IC_50_ of 0.068nM. Here, C05 on the bat and human constant regions were included as a negative control; it had minimal activity against NiV-pps, reinforcing that ReBAs’ binding activity is antigen-specific and depends on its paratope.

## Discussion

Due to the lack of available bat-specific reagents, there is limited experimental evidence on the functions of bat immunoglobulins. To address this gap we have investigated the production of ReBAs and demonstrate robust expression and purification using a standard mammalian cell expression system, and similar *in vitro* activities compared to human IgG1 counterparts in immunological assays. Using a streamlined InFusion based cloning method previously established for recombinant human mAbs^61^, we were able to generate full length ReBAs that presented the variable domains of neutralising mAbs 5B3 and C05. Reducing SDS-PAGE analysis of affinity purified ReBAs confirmed the presence of both Hc of about 50kDa and a Lc of approximately 25kDa (Figures 2A and 2B). These findings are consistent with previous research that reported wild bat IgG Hc at approximately 50kDa and Lc at approximately 25 kDa^30,34,62,63^. Further characterisation with SEC and negTEM found whole ReBAs resembles human IgG1 in terms of oligomeric state, molecular weight and Y-shape feature (Figures 2C, 2D and 2F). Together, these results showed that our expression plasmid could produce functional recombinant bat IgG1 analogous to human IgG1.

In this study, recombinant bat antibodies were successfully purified using pA and pG columns, supporting our hypothesis that ReBAs are mammalian-like and could be recovered using conventional IgG purification methods (Figures 2A and 2B). This finding is consistent with our structural and sequence analysis that found that binding sites for both pA and pG are largely conserved between human and bat IgG1 Fc from both suborder Yinpterochiroptera (*P.alecto)* & Yangochiroptera (*M.brandtii)* (Figure S7)^64–66^. Intriguingly, this contradicts previous findings that found sera-derived bat IgGs from the same Pteropodidae family preferentially bind pG over pA^30,63^. One possible explanation is that bat sera may contain multiple subclasses of IgG with different binding affinities to pA and pG. A similar observation has been reported in other mammalian immunoglobulins and may explain why ReBAs, a bat IgG1, is indifferent to pA or pG purification^67^. Purification with protein L was not attempted as it was previously shown to be ineffective in isolating sera-derived bat IgG from flying foxes despite evidence of kappa Lc usage in some bats^18,30,34^. It would also be interesting for future studies to investigate if this is due to the usage frequency of kappa and lambda genes or divergence in amino acid sequence in the kappa Lc of bats.

Next, we evaluated how ReBAs expression is affected by Hc signal sequences. The signal sequence encodes for a short peptide that guides nascent protein through the secretory pathway^68^. Our results show that signal peptides derived from human antibodies (Hsp) or Bat antibody (Bsp) are largely comparable in the ExpiCHO system (Figure 2C and D). Still, it would be interesting for future work to explore the impact of divergent signal peptides from different bat species and assess protein expression in bat cell line, which may improve expression yield and represents a more native cellular environment for ReBAs expression.

To demonstrate that ReBAs with bat constant region are immunologically functional, we tested them in binding and neutralisation assays. Here, h5B3 showed subnanomolar apparent K_D_ against prefusion NiV F, which is consistent with our previous study^60^. Interestingly, ReBAs purified with pA performed slightly better than those purified with pG in ELISA format (Figures 3 and S5). This disparity could be explained by the fact that pG binds to both Fab & Fc region and Lc plasmid was transfected in excess, potentially resulting in inflated total amount of intact IgG1 by free Lc^69^. Alternatively, the harsher elution conditions (pH 2.7 vs pH 3) required by pG could potentially lead to partial denaturation. This could be remedied in part by a single plasmid system and adopting affinity tags with neutral elution conditions, such as C-tag^70,71^. Additionally, we note that h5B3 and hC05 exhibited higher magnitude of binding (B_max_) to respective antigens than both versions of bat derivatives (Figures 3 and S6). This observation may be skewed by the usage of different species-specific secondary antibodies against human and bat IgG, Therefore, the comparison between human and bat IgG should be interpreted with care. Overall, these findings confirmed that ReBAs recapitulate the specificity and binding capacity to antigens with a bat IgG1 framework.

Our pseudovirus neutralisation assay is consistent with our ELISA, demonstrating that ReBAs exhibit a neutralisation profile comparable to with that of human antibodies with the same paratope. However, our work focuses on 5B3 and C05 and their interactions with their respective antigens. Whether the loss of a single cysteine in the hinge region of bat antibody could impact other paratopes or influence Fc-mediated effector function warrants further investigation. Collectively, in this work, we provide the first framework for the recombinant expression of chimeric bat antibodies that are immunologically functional and bear canonical features of mammalian IgG1. A natural progression of this work is to examine whether ReBAs can mediate Fc-mediated antibody effector functions. Lastly, reagents developed in this study represent an important addition of bat-specific tools for future researchers to understand better how bat antibodies combat viruses.

## Materials and Methods

### Cell Culture

HEK293T cells (ATCC CRL-3216) were cultured in DMEM (Gibco) supplemented with 10% heat-inactivated FCS (Bovogen), 1% of 10,000 U/mL penicillin and streptomycin (Gibco) and 1mM sodium pyruvate at 37°C in a humidified incubator with 5% CO_2_. BHK-21 cells (ATCC CCL-10) were cultured in DMEM (Gibco) supplemented with 5% heat-inactivated fetal calf serum (Bovogen), 1% of 10,000 U/mL penicillin and streptomycin (Gibco) at 37°C in a humidified incubator with 5% CO_2_. ExpiCHO (Thermofisher Scientific) were cultured in ExpiCHO expression medium (Thermofisher Scientific) at 37°C in a humidified incubator with 7.5% CO_2_.

### Recombinant antibody and protein expression and purification

To generate recombinant bat antibodies against henipavirus fusion protein, the variable region of heavy (VH) and light chain (VL) of murine 5B3 or C05 were cloned into bat IgG1 framework region (GenBank: GQ427152.1, ELK10654.1) with human (Hsp) or bat signal peptide (Bsp) sequence in pNBF expression plasmids (National Biologics Facility).^50,54,55,72^ All constructs were sequence verified by Australian Genome Research Facility (AGRF). Detailed information on construct and signal peptides can be found in Figures 1 and S1. Cloning was performed using In-Fusion cloning and stellar competent cells (TakaraBio) as per the manufacturer’s recommendation. In brief, expression plasmids containing 15μg of light and 10μg heavy chains were added to ExpiCHO cells at a density of 6x10^6^ cells per mL. Seven days post-transfection, cell culture supernatant was harvested by sterile filtration and centrifuging at 4800xg for 30mins. Cell culture supernatant was purified with AKTA Start or Pure (Cytiva) using HiTrap Protein A or Protein G antibody purification columns (Cytiva). Protein A column was washed with buffer containing 25mM Tris Base, 25mM NaCl, pH7.4 and eluted with 100mM sodium acetate, 150mM NaCl, pH3. Protein G column was washed with buffer containing 20mM sodium pyruvate, 150mM NaCl, pH7.4 and eluted with 100mM glycine, pH2.7. Antibodies were then sterile filtered, concentrated and buffer exchanged into PBS pH7.4, and concentration quantified using Nanodrop One (ThermoFisher). For antigens, NiV F and Influenza HA (A/Brisbane/59/2007) Clamps were produced as previously described^57,60,73^.

### Protein characterisation by SDS-PAGE and size-exclusion chromatography (SEC)

Using SDS-PAGE (Bio-Rad), molecular weight and purity of proteins were assessed under reducing (100mM dithiothreitol) by loading 5μg of boiled sample onto stacking (4%) and resolving (12.5%) polyacrylamide gel. After gel electrophoresis, the gel was stained with R-250 Coomassie brilliant blue for 1 hour, destained overnight with 40% methanol and rinsed with Milli-Q water. Using analytical SEC, purified antigens and antibodies were further evaluated for aggregates and oligomeric states. 50-100μg of sample in PBS were manually loaded onto a 500μL loop before passing it through Superose 6 Increase 10/300 GL column and AKTA Pure (Cytiva). Samples corresponding to retention volume with absorbance peaks were collected in 1mL fractions in 96-well DeepWell plates and data we normalised to the highest absorbance observed per run as relative mAU. The molecular weight of proteins was established with reference to three standard proteins.

### Enzyme-linked immunosorbent assay (ELISA)

Nunc MaxiSorp flat-bottom plates (ThermoFisher) were coated with 50μL PBS containing antigens at a concentration of 2μg/mL and incubated overnight at 4°C. Nonspecific binding was blocked by incubating all wells on plate with 150μL of blocking agent (5% Milk Diluent (SeraCare) in PBS with 0.1% TWEEN-20 for one hour at room temperature. After blocking, antigens were probed with 50μL of serially diluted sera from immunised bats (starting at 1 in 10 dilution) or primary antibodies (Human: h5B3, hC05, anti-Clamp; Bat: b5B3Hsp, b5B3Bsp, bC05Hsp, bC05Bsp; Goat: goat anti-bat IgG conjugated to HRP (Novus Biologicals)) and incubated at 37°C for one hour. Plates were then washed thrice in water before adding 50μL of secondary antibodies conjugated with HRP (goat anti-bat (Novus Biologicals, Alpha Diagnostic and Bethyl) or goat anti-human (Sigma Aldrich) were used at a concentration of 0.334μg/mL and 0.167μg/mL, respectively). Plates were then incubated at 37°C for one hour and were subsequently washed thrice in tap water before drying on a paper towel. After drying, 50μL of warmed TMB (ThermoFisher) was added to plates and developed at room temperature for 5 minutes. 25μL of 1M H_2_SO_4_ was added to stop the enzymatic reaction before reading the plates at 450nm on Varioskan LUX (ThermoFisher). The background signal was determined by PBS-only wells and subtracted from all readings.

### Negative-stain transmission electron microscopy (negTEM)

SEC purified proteins (10μg/mL) were coated onto glow-discharged carbon-coated grids (EMS) and incubated for 2 minutes. Grids were then rinsed with three drops of Milli-Q water and stained with 2% uranyl acetate. Micrographs of the sample were collected with HITACHI HT7700 operated at 120 keV with 40,000x magnification.

### Glycopeptidase F sensitivity analysis

Ten micrograms of purified antibodies were boiled for 10 minutes and chilled on ice. Samples were then reduced with 1μL NP-40, and N-linked oligosaccharides were removed by adding 1μL of PNGase F enzyme (New England Biolabs) before 1 hour incubation at 37°C. The glycosylation state of samples was then analysed with PAGE as described in SDS-PAGE protocol above.

### Pseudovirus neutralisation assays

Pseudovirus neutralisation assays were performed using lentivirus-based pseudotypes as previously described^58,60^. Briefly, HEK293T cells were transfected with p8.91 (encoding for HIV-1 gag-pol), CSFLW (lentivirus backbone expressing a firefly luciferase reporter gene) and viral glycoproteins (NiV F and G) using LTX transfection reagent. Supernatants containing pseudotyped virus were harvested at 48- and 72-hours post-transfection, pooled and centrifuged at 1,300x*g* for 10 minutes at 4°C to remove cellular debris. For the neutralisation test, HEK293T cells were then seeded overnight at a density of 2 ×10^4^ in 100µL and incubated at 37°C, 5% CO_2_. The antibodies were diluted in serum-free media in triplicate titrated 5-fold and incubated with pseudo-particles added at a dilution equivalent to 10^6^ signal luciferase units in DMEM-10% to final volume of 100µL. The complex was incubated for 1 hour at 37°C, 5% CO_2_. Firefly luciferase activity was then measured with BrightGlo luciferase reagent and a Glomax-Multi^+^ Detection System (Promega) after 48 hours post infection. Pseudotyped virus neutralisation titres were calculated by interpolating the point at which there was 50% reduction in luciferase activity, relative to control antibody.

### Bat Immunisation

Immunisations were performed at Colorado State University using a closed, specific pathogen free colony of Jamaican fruit bats. All animal procedures were approved by the Colorado State University (CSU) Institutional Animal Care and Use Committee (protocol 1085) and were in compliance with U.S. Animal Welfare Act. CSU has a captive colony of Jamaican fruit bats, a neotropical fruit bat indigenous to much of South America, Central America and the Caribbean. Colony bats are kept in a free flight room measuring 19’w x 10’l x 9’h. Roosting baskets are hung from the ceiling throughout the room, and drapes of different cloth materials are positioned for hanging and roosting. Ambient temperature is maintained between 20°C and 25°C, humidity between 50% and 70%, and a 12-hour light/12-hour dark-light cycle via a computer-controlled system. Diets consist of a combination of fruits (Shamrock Foods, Fort Collins, CO), Tekald primate diet (Envigo, Huntington, UK), molasses, nonfat dry milk and cherry gelatin that are placed in multiple feeding trays around the room once a day. Fresh water is provided. In addition, fruit is hung around the room to stimulate foraging behaviour and serve as enrichment. Bats were immunised twice with NiV F proteins adjuvanted with Addavax (InvivoGen) before blood collection 14 days apart. Blood was collected 14 days after the booster. A maximum blood volume between 1 and 1.5mL is collected in a syringe and transferred to a red top tube (RTT). RTTs sat at room temperature for one hour to allow a clot to form and then centrifuged at 1000xg for 10 min at room temperature. Serum was removed from the clot, placed in a new microcentrifuge tube and stored at -20°C.

Figures

Supplementary Materials

#Ref to attached illustrator images

## Acknowledgments

The authors would like to thank Dr Christopher McMillan (School of Chemistry and Molecular Biosciences) for kindly providing the haemagglutinin antigen and hC05 antibodies used in this study.

## Author contributions

J.H., A.I., N.M., and D.W. conceptualised the experiment. J.H., A.I., B.L., T.S., N.M., and D.W. contributed to experiments. T.S., K.J.C., P.R.Y., N.M., and D.W. provided resources. J.H., A.I, N.M., and D.W. performed data analysis. J.H. wrote the draft of the manuscript. J.H. A.I., N.M., and D.W. edited the manuscript. K.J.C., P.R.Y., N.M., and D.W. acquired funding. N.M., and D.W. supervised the work.

---

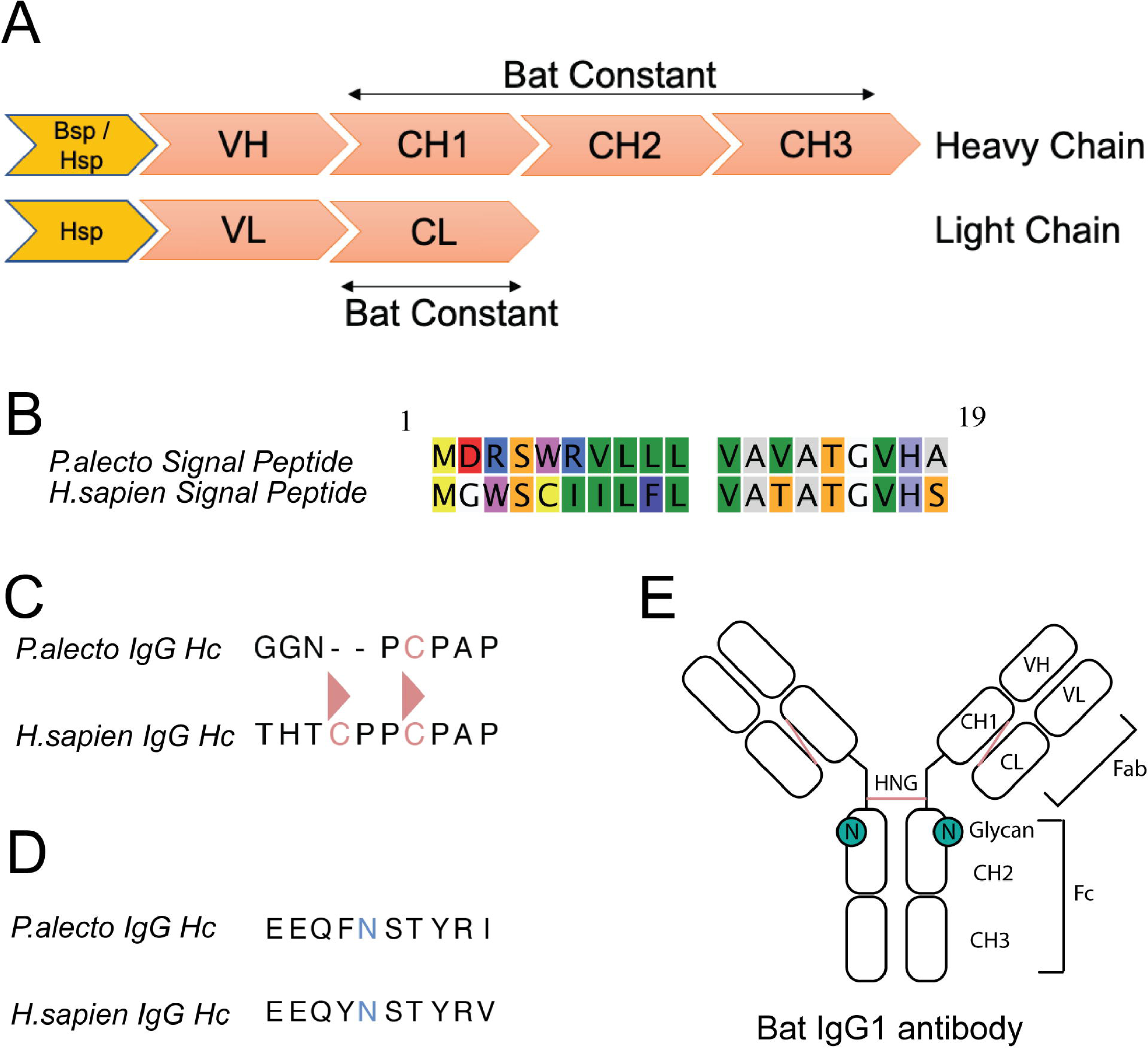

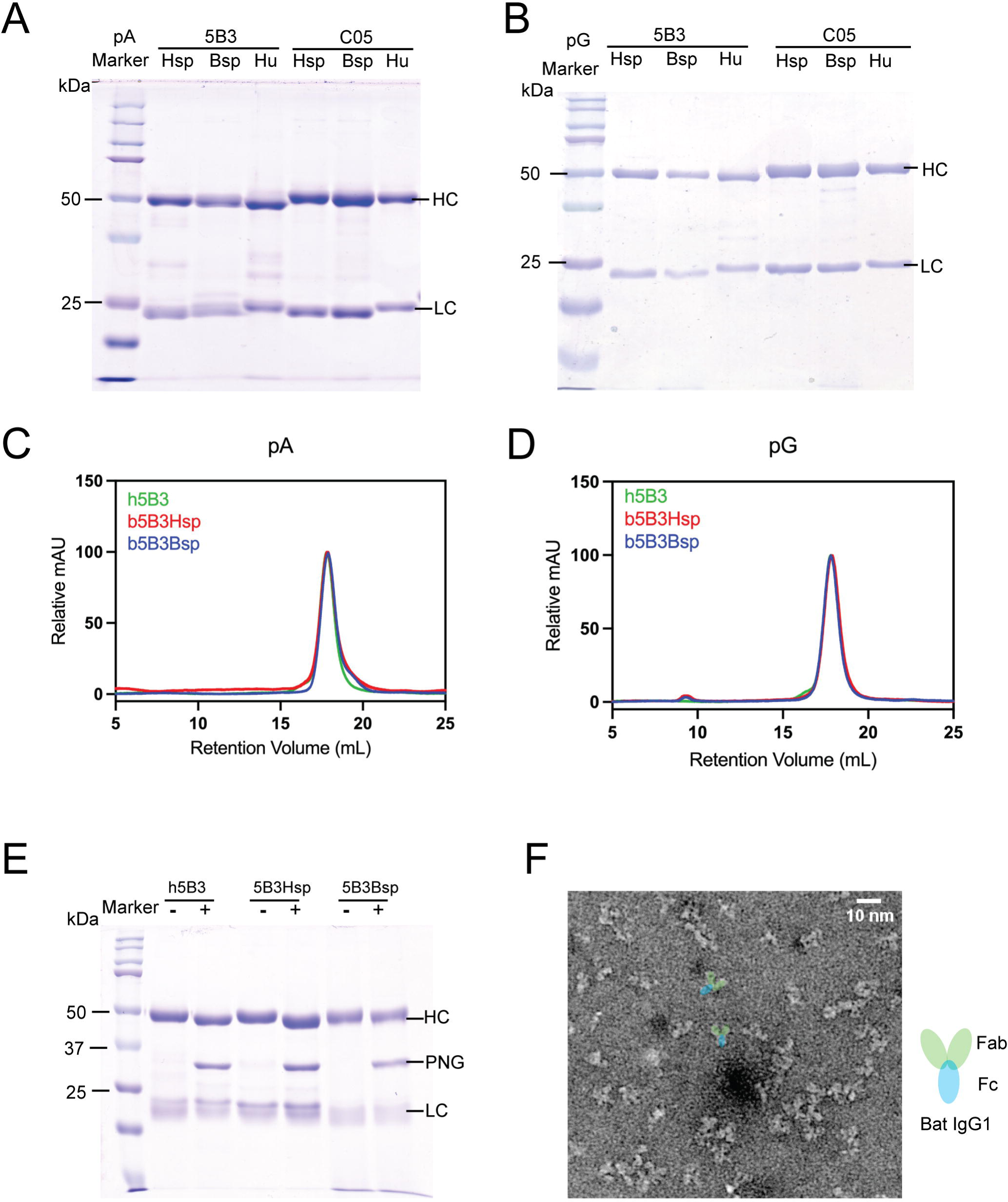

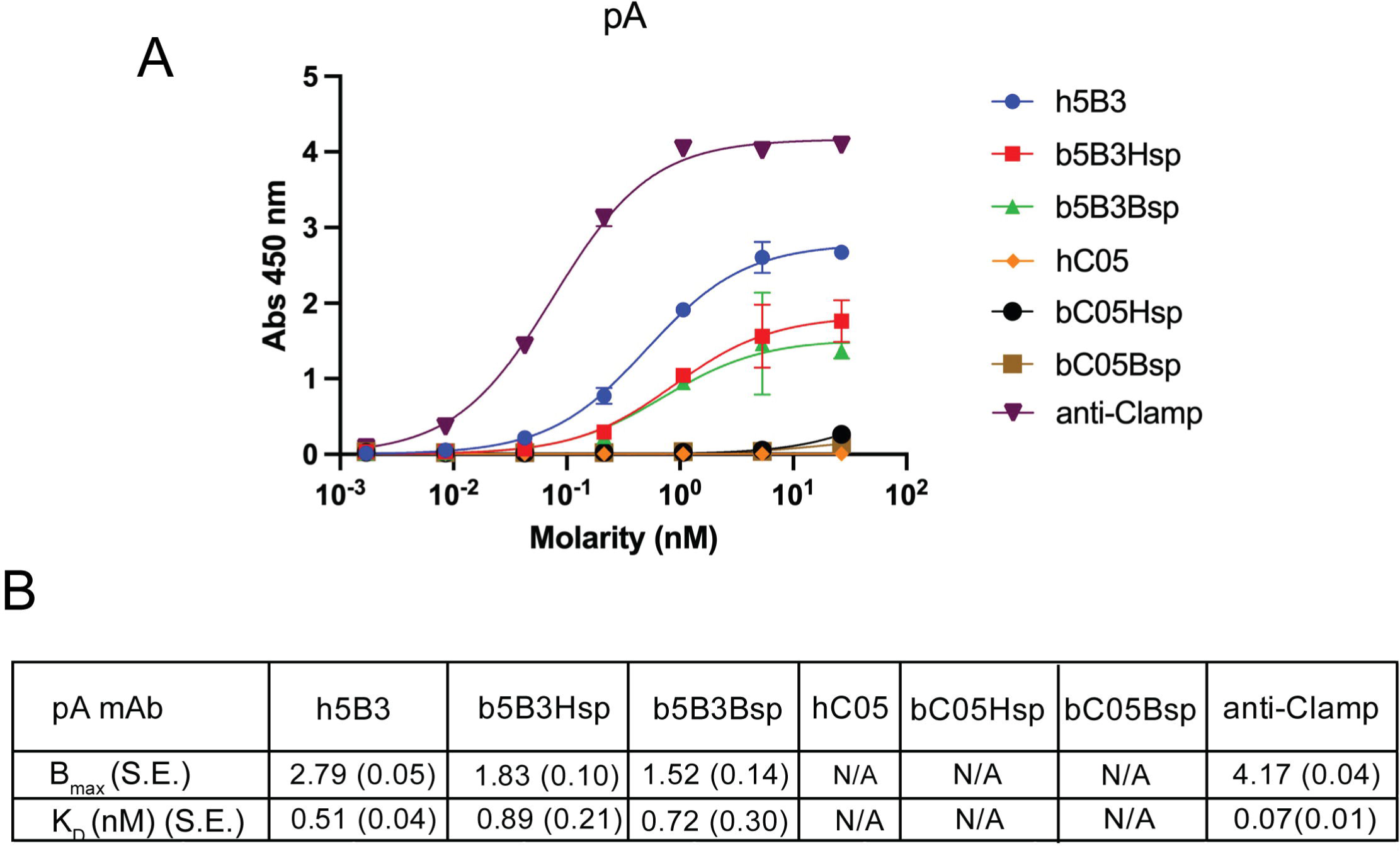

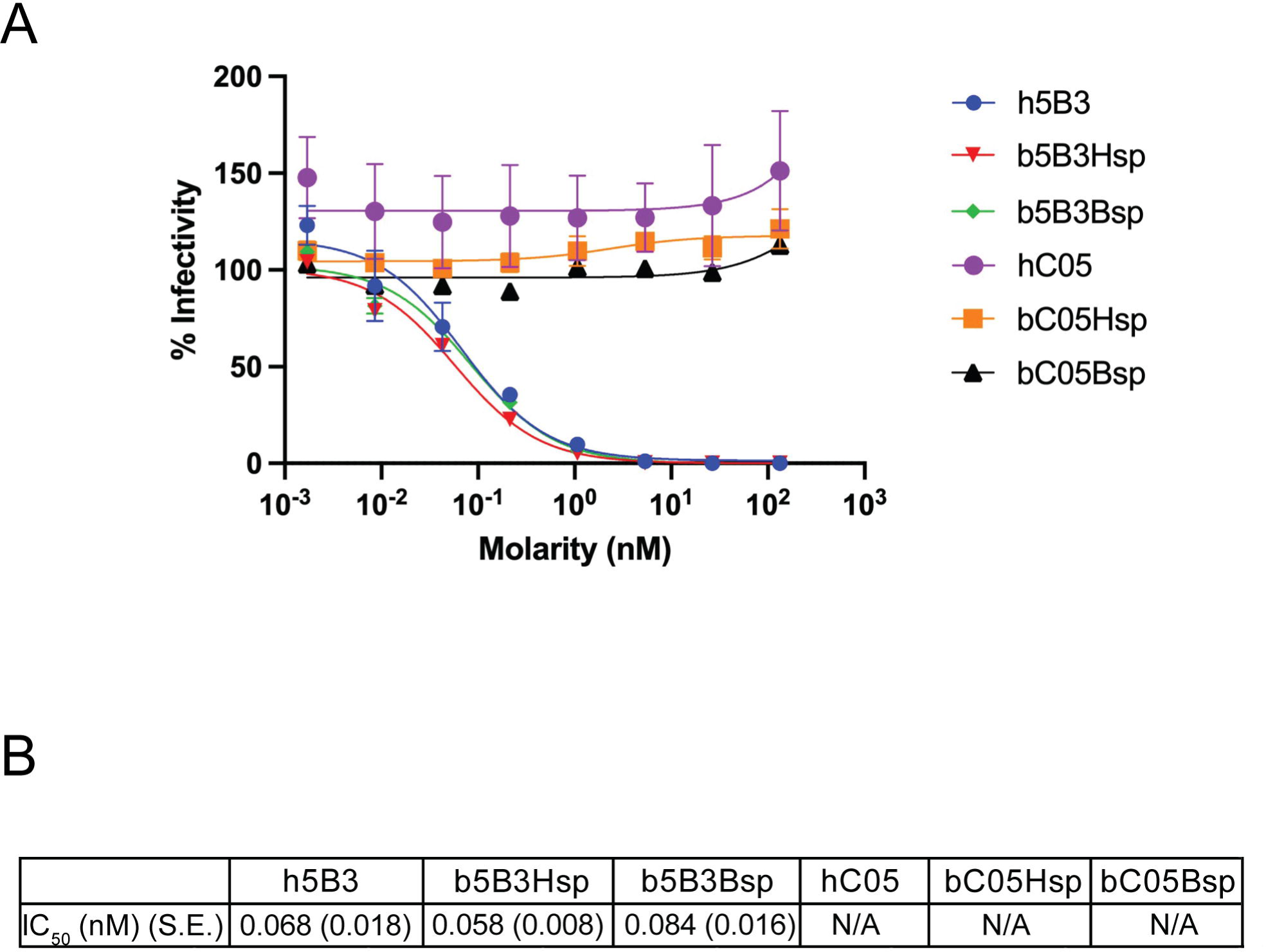

