## supplementary for "Cloning, expression and characterisation of antigen-specific recombinant bat immunoglobulin from the black flying fox (*Pteropus alecto*)"

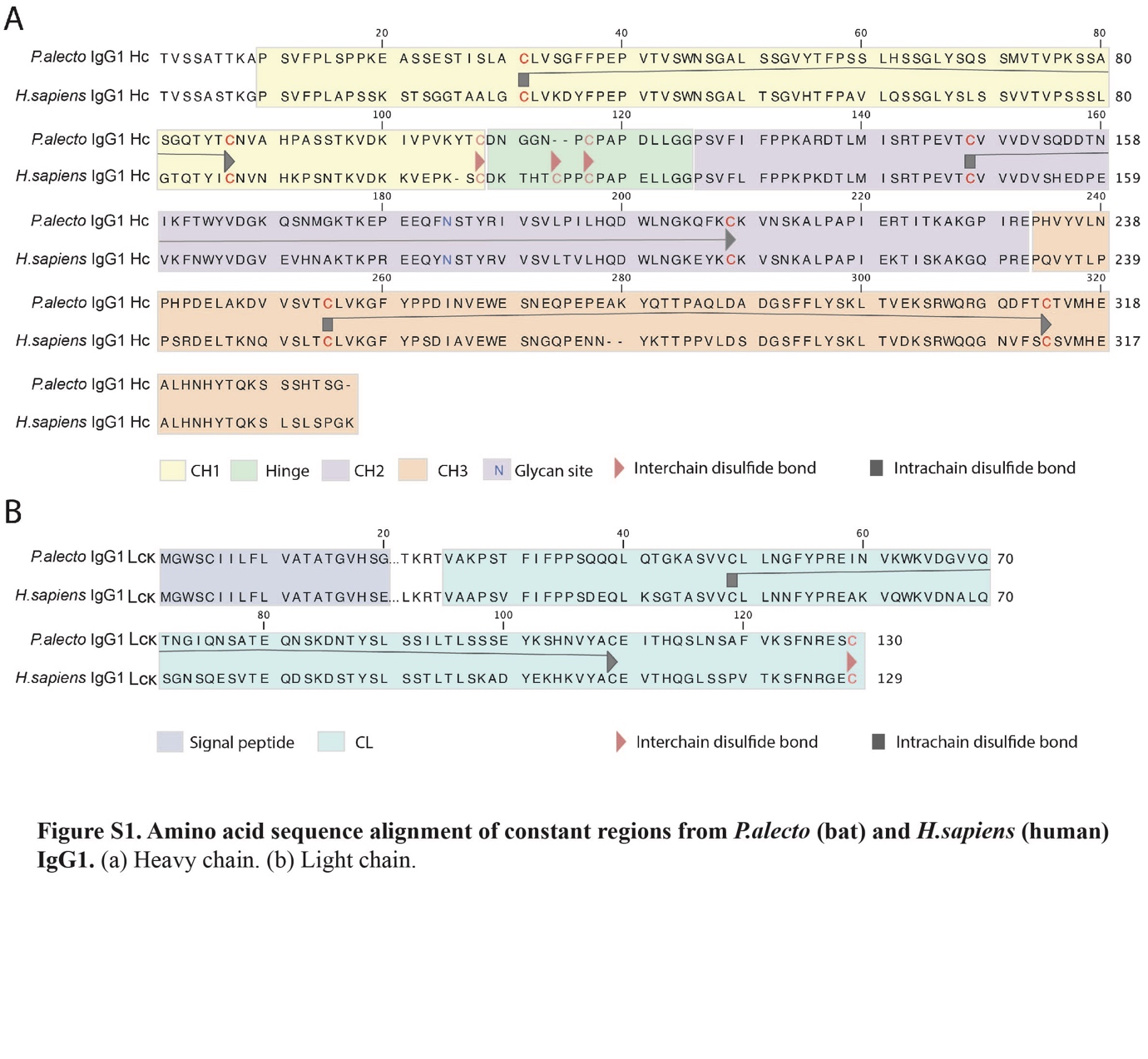

**Figure S1. Amino acid sequence alignment of constant regions from *P.alecto* (bat) and *H.sapiens* (human) IgG1.** (a) Heavy chain. (b) Kappa Light chain.

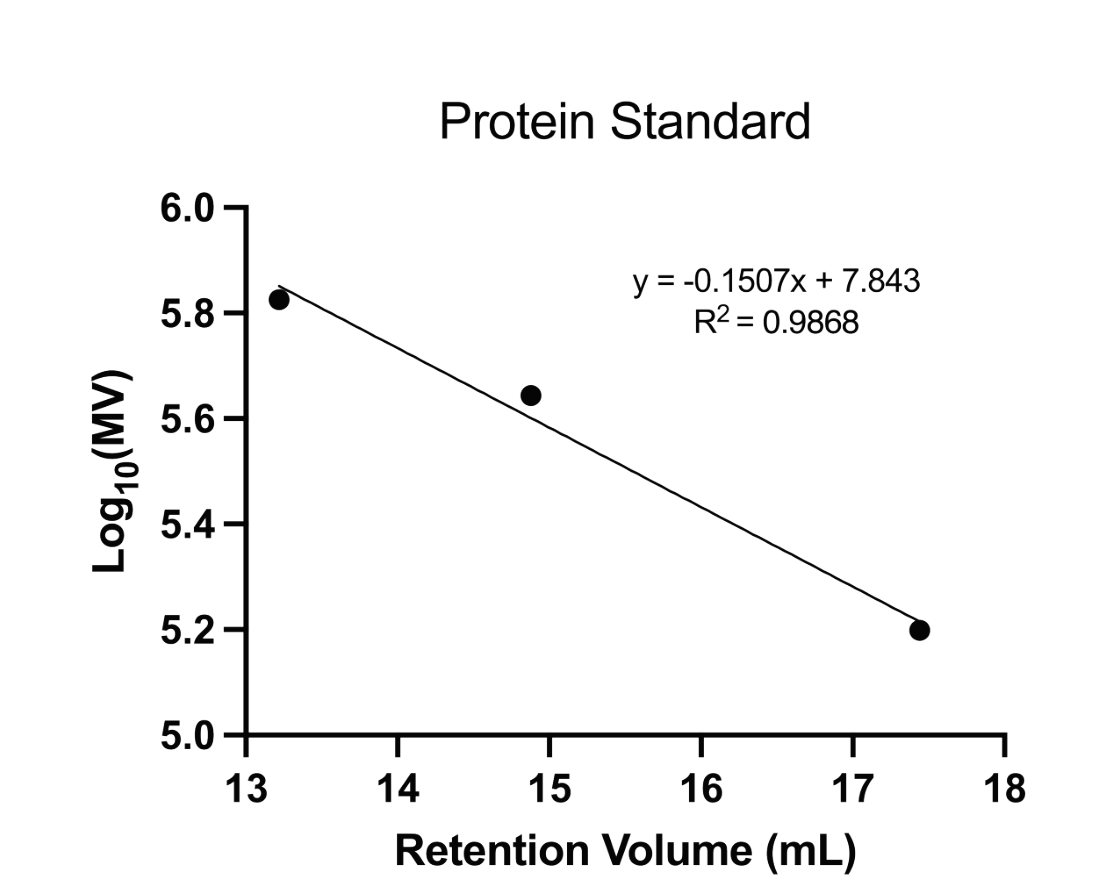

**Figure S2. Chromatogram depicting size exclusion analysis of three standard protein with known molecular weight (Thyroglobulin, 699kDa; Ferritin, 440kDa; and Aldolase, 158kDa).** A linear regression model was generated using these three standard proteins. Substitution of the ReBAs elution peak retention volume into this model then yield an estimated molecular weight of approximately 145 kDa for ReBAs.

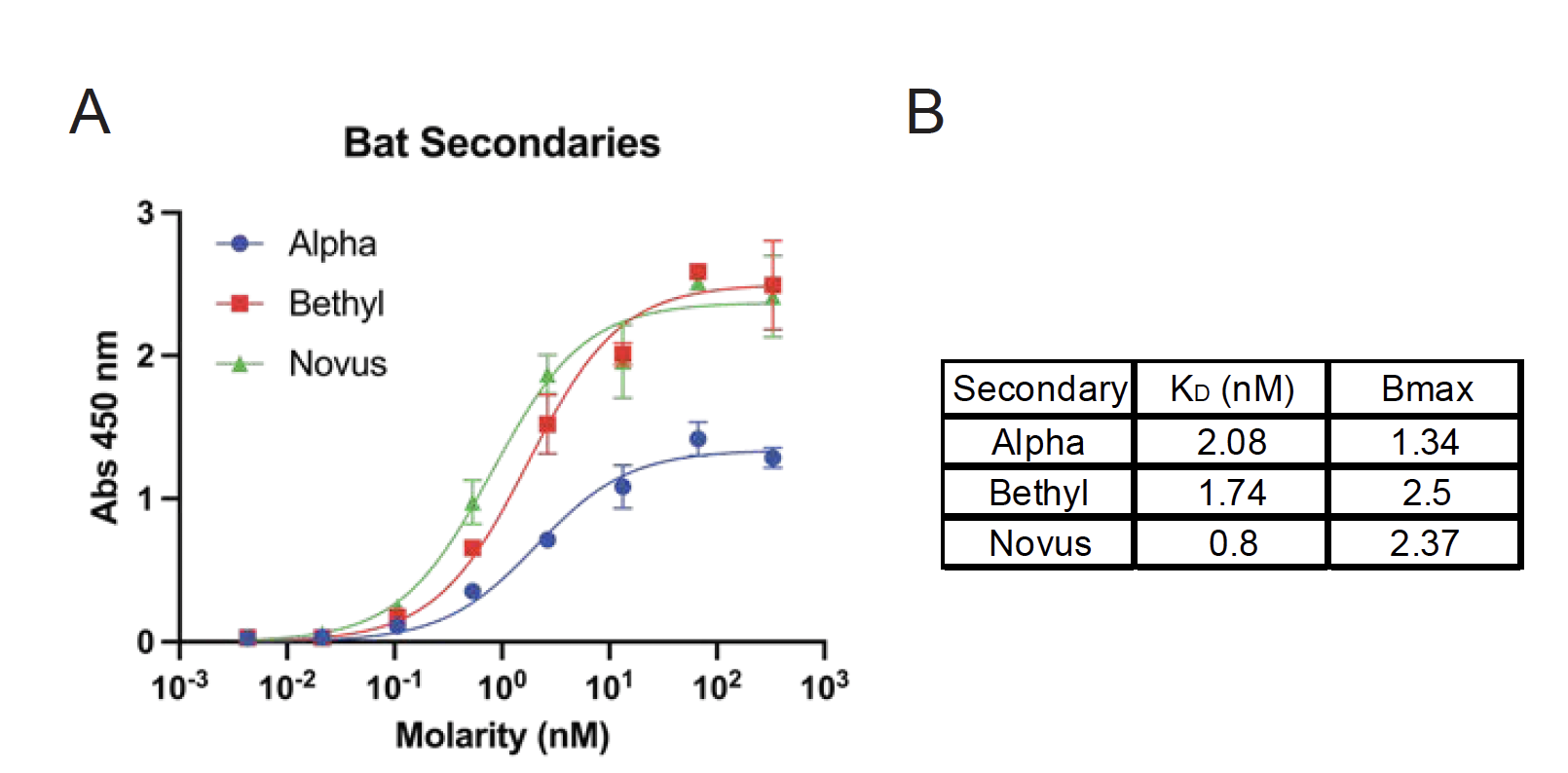

**Figure S3. Validation of commercial HRP-conjugated secondary antibodies against ReBA.** (a) Indirect ELISA absorbance readings of three commercial secondary antibodies directed against ReBAs. (b) K_D_ in nM & B_max_ indicates that Novus Goat anti-bat IgG (Cat No: NB7238) has the highest affinity to ReBA derived from *P.alecto* (Black flying fox). Data presented here represents two technical replicates

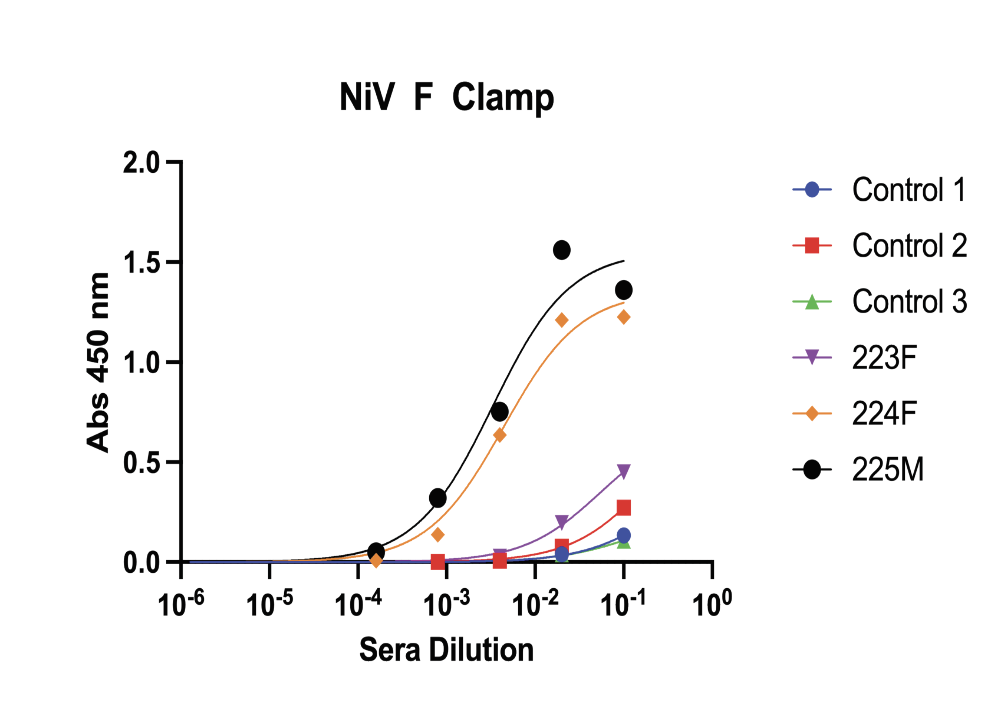

**Figure S4. Indirect ELISA shows that 2 of 3 *A.jamaicensis* (Jamaician fruit bat) immunised with NiV F antigens seroconverted against prefusion NiV F antigen stabilised by Clamp trimer.** Goat anti-bat IgG conjugated with HRP (Cat No: NB7238) is utilised as secondary. Data presented here represents two technical replicates.

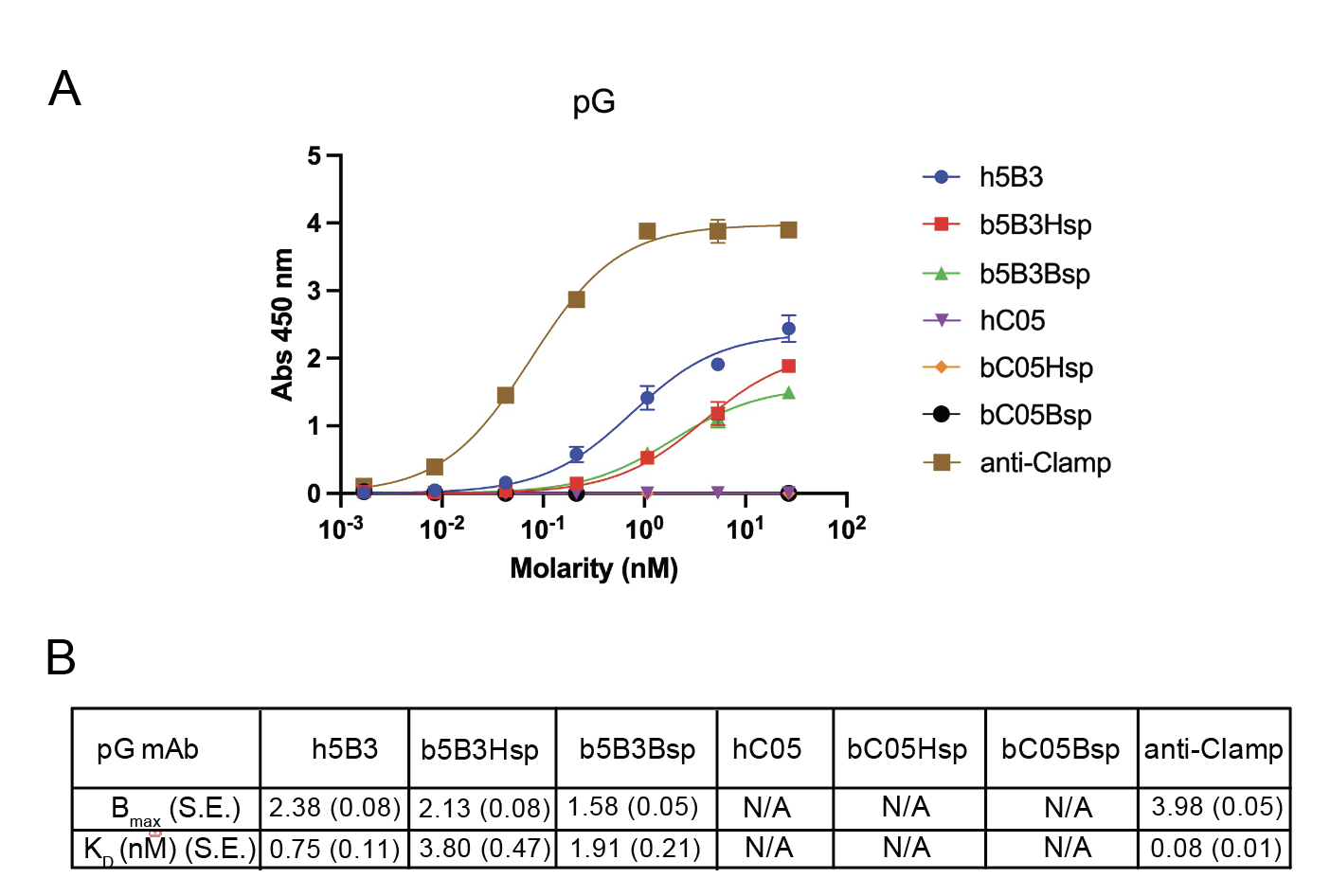

**Figure S5. Recombinant bat 5B3 antibodies binds target antigen in indirect ELISA format.** (a) Indirect ELISA absorbance readings of ReBAs and human mAb purified with pG directed against prefusion NiV F Clamp antigen. Anti-Clamp mAb (HIV1281) and C05 mAbs are included here as controls. HRP conjugated secondaries are specific to species of primary antibodies. (b) Indirect ELISA B_max_ and K_D_ in nM of ReBAs purified with pG. Data presented here represents two technical replicates. Error bars represent standard deviation and standard error (S.E.) is shown in parentheses. N/A; Not Applicable.

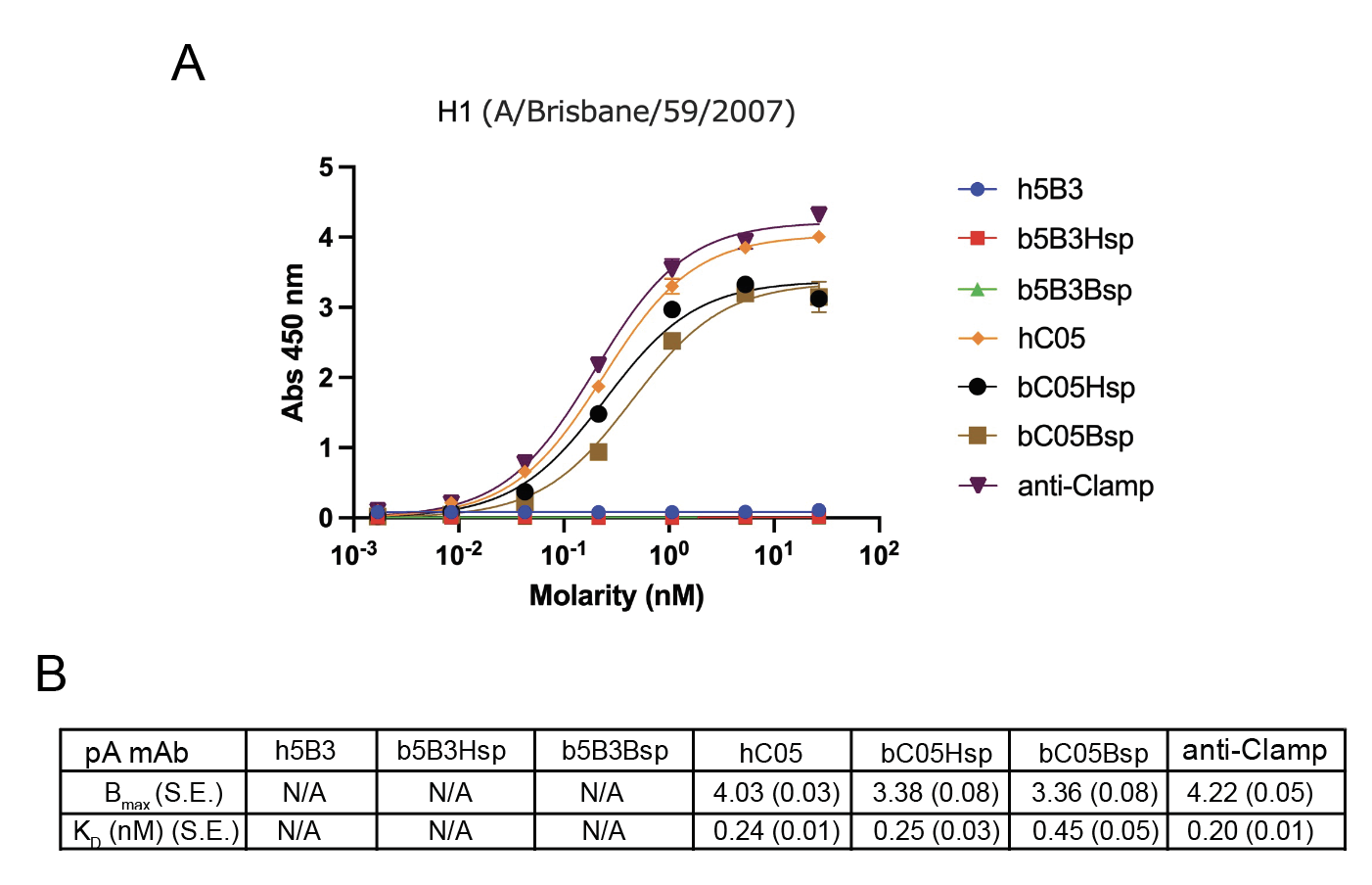

**Figure S6. Recombinant bat C05 antibodies binds target antigen in indirect ELISA format.** (a) Indirect ELISA absorbance readings of ReBAs and human mAb purified with pA directed against Influenza H1 (A/Brisbane/59/2007) Clamp. Anti-Clamp mAb (HIV1281) and 5B3 mAbs are included here as controls. HRP-conjugated secondaries are specific to species of primary antibodies. (b) Indirect ELISA B_max_ and K_D_ in nM of ReBAs. Data presented here represents two technical replicates. Error bars represent standard deviation and standard error (S.E.) is shown in parentheses. N/A; Not Applicable.

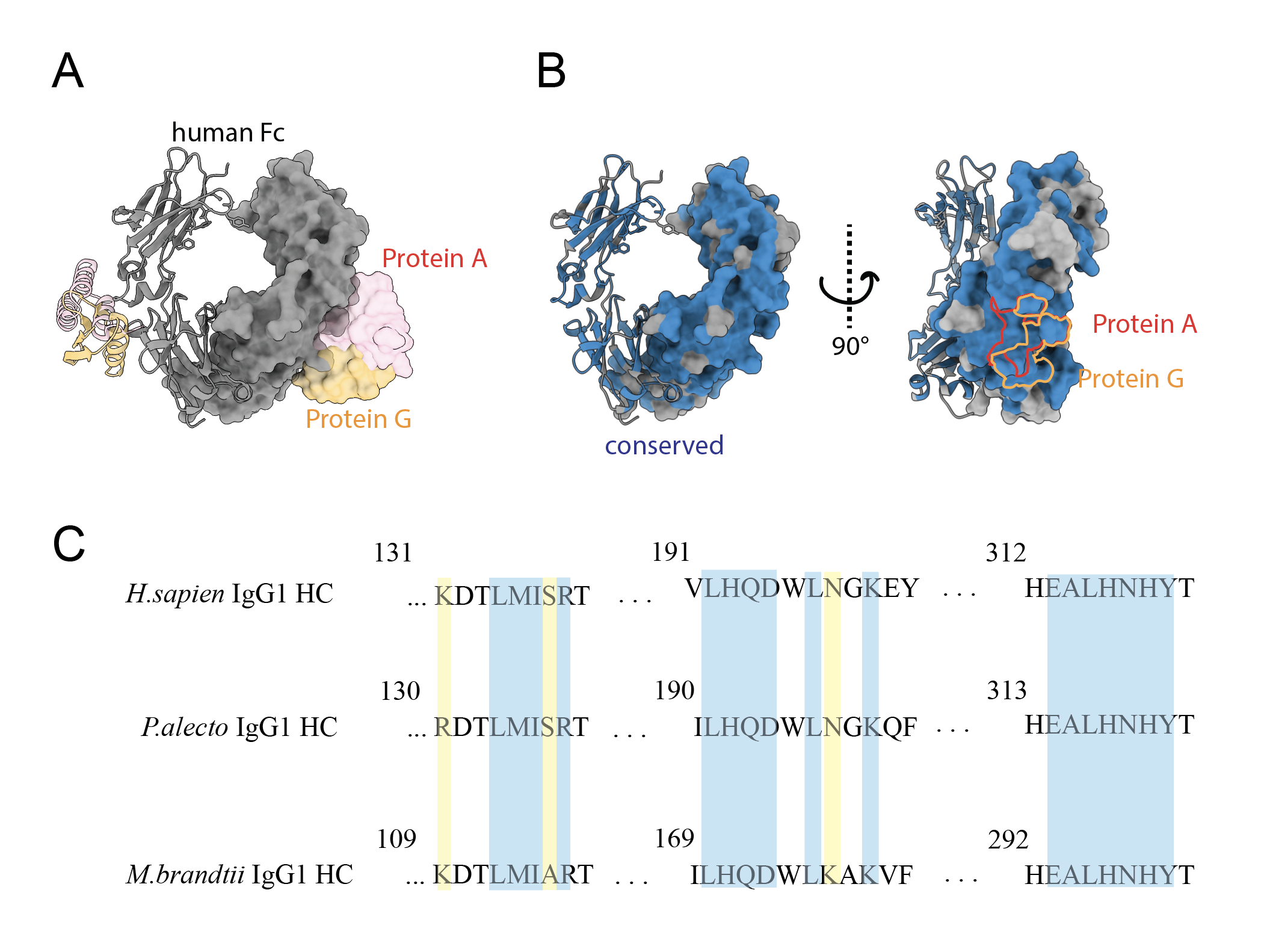

**Figure S7. Comparison of footprint of Protein A and Protein G on Fc region of *H.sapiens* (human) and *P.alecto* (bat) IgG1.** (a) Molecular surface representation of human Fc showing in grey, engaging with Protein A showing in pink and Protein G showing in orange. (b) Conserved residues of bat Fc showing in blue on molecular surface representation of human Fc showing in grey. Footprint of Protein A highlighted in red and Protein G highlighted in orange indicates that both would bind bat Fc region. (c) HC sequence alignment of *H.sapiens*, *P.alecto (*ADD71697.1), and *M.brandtii* (EPQ16374.1) shows that majority of Protein A binding residues are conserved (blue) with exceptions shown in yellow. Crystal structure of Protein A and Protein G binding site is modified from 4WWI and 1FCC respectively.
